# Model-based prediction of muscarinic receptor function from auditory mismatch negativity responses

**DOI:** 10.1101/2020.06.08.139550

**Authors:** Dario Schöbi, Fabienne Jung, Stefan Frässle, Heike Endepols, Rosalyn J. Moran, Karl J. Friston, Marc Tittgemeyer, Jakob Heinzle, Klaas Enno Stephan

## Abstract

Drugs affecting neuromodulation, for example by dopamine or acetylcholine, take centre stage among therapeutic strategies in psychiatry. These neuromodulators can change both neuronal gain and synaptic plasticity and therefore affect electrophysiological measures. An important goal for clinical diagnostics is to exploit this effect in the reverse direction, i.e., to infer the status of specific neuromodulatory systems from electrophysiological measures.

In this study, we provide proof-of-concept that the functional status of cholinergic (specifically muscarinic) receptors can be inferred from electrophysiological data using generative (dynamic causal) models. To this end, we used epidural EEG recordings over two auditory cortical regions during a mismatch negativity (MMN) paradigm in rats. All animals were treated, across sessions, with muscarinic receptor agonists and antagonists at different doses. Together with a placebo condition, this resulted in five levels of muscarinic receptor status. Using a dynamic causal model - embodying a small network of coupled cortical microcircuits - we estimated synaptic parameters and their change across pharmacological conditions. The ensuing parameter estimates associated with (the neuromodulation of) synaptic efficacy showed both graded muscarinic effects and predictive validity between agonistic and antagonistic pharmacological conditions.

This finding illustrates the potential utility of generative models of electrophysiological data as computational assays of muscarinic function. In application to EEG data of patients from heterogeneous spectrum diseases, e.g. schizophrenia, such models might help identify subgroups of patients that respond differentially to cholinergic treatments.

**Significance Statement:** In psychiatry, the vast majority of pharmacological treatments affect actions of neuromodulatory transmitters, e.g. dopamine or acetylcholine. As treatment is largely trial-and-error based, one of the goals for computational psychiatry is to construct mathematical models that can serve as “computational assays” and infer the status of specific neuromodulatory systems in individual patients. This neuro-modelling strategy has great promise for electrophysiological data in particular but requires careful validation. The present study demonstrates that the functional status of cholinergic (muscarinic) receptors can be inferred from electrophysiological data using dynamic causal models of neural circuits. While accuracy needs to be enhanced and our results must be replicated in larger samples, our current results provide proof-of-concept for computational assays of muscarinic function using EEG.

## Introduction

Many pathophysiological theories of psychiatric conditions emphasize abnormalities of neuromodulation through transmitters such as dopamine or acetylcholine (Tandon and Greden, 1989; Cohen and Servan-Schreiber, 1992; Stephan et al., 2006; Howes and Kapur, 2009; Higley and Picciotto, 2014). Indeed, most drugs used in clinical psychiatry affect synthesis, reuptake, or postsynaptic action of neuromodulatory transmitters. However, patients with the same diagnosis according to ICD/DSM often show great variability in their response to the same treatment, a likely consequence of pathophysiological heterogeneity under contemporary diagnostic classification schemes (Kapur et al., 2012; Krystal and State, 2014; Stephan et al., 2016). There is thus a pressing need for clinical tests that pinpoint specific abnormalities of neuromodulation in individual patients.

The present study is motivated by mechanistic theories that highlight cholinergic abnormalities in schizophrenia (Friston, 1998; Stephan et al., 2006; Stephan et al., 2009a) with a specific focus on muscarinic receptors (Raedler et al., 2007; Scarr and Dean, 2008). Empirically, both *post-mortem* and *in vivo* studies have provided evidence for abnormalities in muscarinic receptor availability (Raedler et al., 2003; Scarr et al., 2009; Scarr et al., 2013). Importantly, a ‘muscarinic receptor-deficit schizophrenia’ (MRDS) subgroup was identified that was unrelated to treatment, illness duration, gender or age and characterized by substantially decreased numbers of muscarinic receptors in dorsolateral prefrontal cortex and associated differences in gene expression and synaptic properties (Scarr et al., 2009; Gibbons et al., 2013; Scarr et al., 2013; Dean et al., 2015; Scarr et al., 2018). These marked differences in muscarinic receptor function – across the schizophrenia spectrum – have implications for treatment: not least because clozapine and olanzapine, two antipsychotics with particular efficacy but also side effects, have distinctive antagonistic activity at muscarinic receptors (Weiner et al., 2004; Raedler, 2007) (for a comparative overview of antipsychotics, see (Kapur and Remington, 2001)). Therefore, if muscarinic receptor status could be determined non-invasively and cost-efficiently in individual patients, this might guide personalized treatment selection.

Unfortunately, with the exception of specialized positron emission tomography procedures, we currently lack non-invasive *in vivo* measures of neuromodulatory transmitters in humans. An alternative approach rests on generative models as computational assays of neuromodulation (Stephan et al., 2006; Friston et al., 2013; Stephan and Mathys, 2014). For example, dynamic causal models (DCM; David et al. 2006) describe how latent neuronal processes generate electrophysiological measures in terms of synaptic parameters that are sensitive to dopaminergic (Moran et al., 2011) and cholinergic alterations (Moran et al., 2013). So far, however, validation studies are lacking that employ more than one kind of pharmacological perturbation and which examine the model’s ability to predict neuromodulatory status out-of-sample.

In this proof-of-concept study, we tested the feasibility of using DCM to infer and predict muscarinic receptor function. We used a rodent model where pharmacological interventions can be repeated in the same animal with different doses and drugs. As an experimental paradigm, we chose the auditory mismatch negativity (MMN) which is reliably impaired in schizophrenia (Baldeweg et al., 2004; Umbricht and Krljes, 2005; Erickson et al., 2016) and is sensitive to cholinergic manipulations (for review, see (Garrido et al., 2009)). Epidural EEG recordings were obtained bilaterally from primary and secondary auditory areas of awake rats, thus avoiding any confounds by anaesthesia. All animals underwent five pharmacological conditions: (i) two dosages of the muscarinic antagonist scopolamine, (ii) vehicle, and (iii) two dosages of the muscarinic agonist pilocarpine. The measured EEG activity was modelled as arising from the neuronal dynamics within a set of connected cortical microcircuits. The animal-specific parameter estimates of this generative circuit model served as features for subsequent out-of-sample predictions (i.e., ‘generative embedding’; (Brodersen et al., 2011)).

This approach allowed us to test whether dose-dependent changes in muscarinic receptor function could be predicted, based on estimates of neuronal processes in cortical circuits, from EEG measurements of individual animals. Permutation statistics on classification accuracies ensured that even in our relatively small sample, the conclusions are protected against overfitting (Varoquaux, 2018).

## Methods

### Data Acquisition

The data for this study were acquired at the Max-Planck-Institute for Metabolism Research at Cologne, Germany. All procedures were approved by the local governmental and veterinary authorities of Cologne (file number 9.93.2.10.35.07.056) and followed ARRIVE standards (Kilkenny et al., 2010). For a detailed description of the acquisition protocol, see (Jung, 2013). In brief, electrodes were implanted over the primary auditory cortex (A1, coordinates relative to bregma: 4 mm posterior, ±8 mm lateral, 4 mm ventral) and posterior auditory field (PAF; secondary auditory cortex; coordinates relative to bregma: 6 mm posterior, ±8 mm lateral, 4 mm ventral) in both hemispheres of ten black hooded rats. Electrode position was determined based on stereotaxic location using the coordinates proposed by Doron et al. (2002), who distinguished the two tonotopic regions based on the firing pattern of single neuron recordings (Doron et al., 2002). Following surgery, animals recovered for ten days. In five sessions, rats received different intraperitoneal injections: 1 or 2 mg/kg of the non-selective muscarinic antagonist scopolamine, 3 or 6 mg/kg of the non-selective muscarinic agonist pilocarpine, or a 0.6 % NaCl-solution (vehicle). In order to avoid interactions between treatments, drug injections were administered every third day, in a counterbalanced order across rats.

Acoustic stimuli were delivered in a sound-attenuated cage using a Tucker Davies Technologies® (TDT, Alachua, USA) System 3 and two free-field magnetic speakers (FF1, TDT). Stimuli consisted of short bursts of band-pass filtered noise with bandwidths between 7-9 kHz and 16-18 kHz, respectively. In total, 1000 tones were presented at a frequency of 2 Hz with 10 % deviant probability. Both bandwidths were used as the standard tone once, in two individual sessions per drug condition. All electrophysiological measures were pre-amplified and transmitted (wireless) to a high frequency receiver (TSE Systems GmbH, Bad Homburg, Germany). This setup allowed the rats to move inside the cage without constraint.

### Non-pharmacological dataset

An additional set of six animals received a placebo only treatment (non-pharmacological dataset). These additional data served to optimize the settings for the subsequent analysis of the pharmacological data. Importantly, they did not enter the main analysis. In brief, the analysis of the non-pharmacological dataset informed the selection of the time window for the classical and model-based (DCM) analysis of the data. Empirical priors for the dynamic causal modelling of pharmacological data were informed by posterior estimates from the non-pharmacological dataset. To this end, we inverted all models for all rats of the non-pharmacological group (individually for the valid hemispheres). We then defined the prior mean *E*[*π*(*θ*)] for the pharmacological group as the average of the posterior means *E*[*p*(*θ*|*y*)] of the non-pharmacological group (averaged over models, rats and hemispheres). As an exception, we retained the default prior for modulatory influences, because these influences may depend upon the pharmacological context. There are two complementary interpretations of the ensuing empirical priors. One can think of them as a Bayesian model average with equal posterior model probabilities.

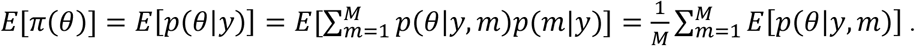

Note that we use *y* as a shorthand notation for the data over all animals and hemispheres.

Alternatively, this corresponds to a parameter average in the space spanned by the parameters that are common to all models. As for the prior variance, we used the variance of posterior means over rats, models, and hemispheres

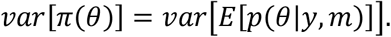

This means that the prior acknowledges the expected between-rat variability.

### Analysis Plan, Data and Code availability

After analysis of the non-pharmacological data but prior to the analysis of the pharmacological data, a version-controlled and time-stamped analysis plan was created. This plan detailed the analysis pipeline ex ante (see Methods section). The analysis plan is provided online at (https://gitlab.ethz.ch/tnu/analysis-plans/schoebietal_auditory_mmn_dcm_2020). Upon acceptance of the paper, the data will be made available in a form adhering to the FAIR (Findable, Accessible, Interoperable, and *Re*-usable) data principles. Furthermore, all analysis is publicly available on https://gitlab.ethz.ch/tnu/code/schoebietal_auditory_mmn_dcm_2020.

### Preprocessing

Preprocessing was implemented using Statistical Parametric Mapping SPM12 (ver. 6906) (Litvak et al., 2011). Electrophysiological data were down-sampled to 1000 Hz (including an anti-aliasing filter), and band-pass filtered between 1 Hz and 30 Hz. Trials exceeding an amplitude of 500 µV were considered artefactual and excluded from further analysis. This (liberal) threshold was chosen based on visual inspection of the single trial ERPs. Comparing the average ERPs before and after artefact rejection showed negligible effects on the averaged waveforms. Finally, standard and deviant tone responses were averaged in a time window of 0 - 250 ms. Following standard procedures for MMN, we averaged standards and deviants, respectively, over all corresponding trials from both sessions, thus removing any potential confounds due to frequency differences in standards and deviants.

All analyses were done individually for each hemisphere. Data from a given hemisphere were excluded if the recording in one of the channels (A1 or PAF) was considered faulty (assessed through visual inspection of the average ERPs prior to any statistical and model-based analyses). This led to exclusion of one left hemisphere and three right hemisphere recordings. We excluded a hemisphere for all pharmacological conditions, even if the recording was of poor quality in only one pharmacological condition.

### Classical Analysis

The MMN paradigm followed a classical oddball design, where the definition of the ‘Standard’ tone frequency did not change throughout a session. We compared the full ERP time series (0 – 250) ms for each of the four electrodes in a fully factorial 2×5×N mixed effects ANOVA (N=9 for left, N=7 for the right hemisphere), with fixed effects factors TONE^1^=[Standard, Deviant], PHARMA=[2mg scopolamine, 1mg scopolamine, vehicle, 3mg pilocarpine, 6mg pilocarpine] and their interaction, and ANIMAL as a random effect (indicated by the notation ‘1|’):

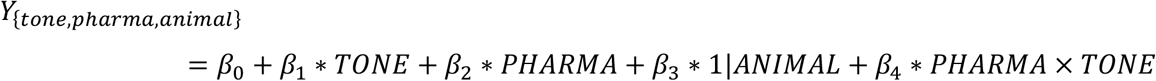

In brief, we fitted the mixed effects ANOVA to every time point in the time window [0, 250] ms post stimulus, where we expected the MMN and thus potential drug effects. The selection of the time window was based on the provisional analysis of the non-pharmacological dataset. We corrected for multiple comparisons using an FDR correction (over time) (Benjamini and Yekutieli, 2001). The analysis only included those rats in which all recordings (in all pharmacological conditions) were valid. Electrodes were analysed separately. We did not additionally correct for electrodes but report the (FDR corrected) p-values to allow for a visual intuition about the significance of the results (e.g. significance threshold under additional Bonferroni correction for p < 0.0125).

All statistical analyses were performed using the open source statistical software R (version 3.5.2) and the packages *lme4, R*.*matlab* and *lmerTest*.

### Dynamic causal modelling

We modelled the data using a convolution based DCM for electrophysiological data (David et al., 2006; Kiebel et al., 2009). In this neural mass model, the average presynaptic firing rate (of a neural population) is transformed into a postsynaptic potential by a convolution operator, while the average potential is converted into average firing rate via a sigmoid activation function. Anatomically, we used a canonical microcircuit (CMC) model (Bastos et al., 2012), where each cortical column (source) comprises two types of pyramidal cell populations, an inhibitory interneuron and an excitatory (spiny stellate) population (Figure 1A). The CMC naturally maps onto computations required for predictive coding (Bastos et al., 2012), which provides a unifying framework for the neuronal computations underlying the MMN and accommodates cortical hierarchies, such as our two-level model (A1 and PAF) (Lieder et al., 2013).

**Figure 1:**
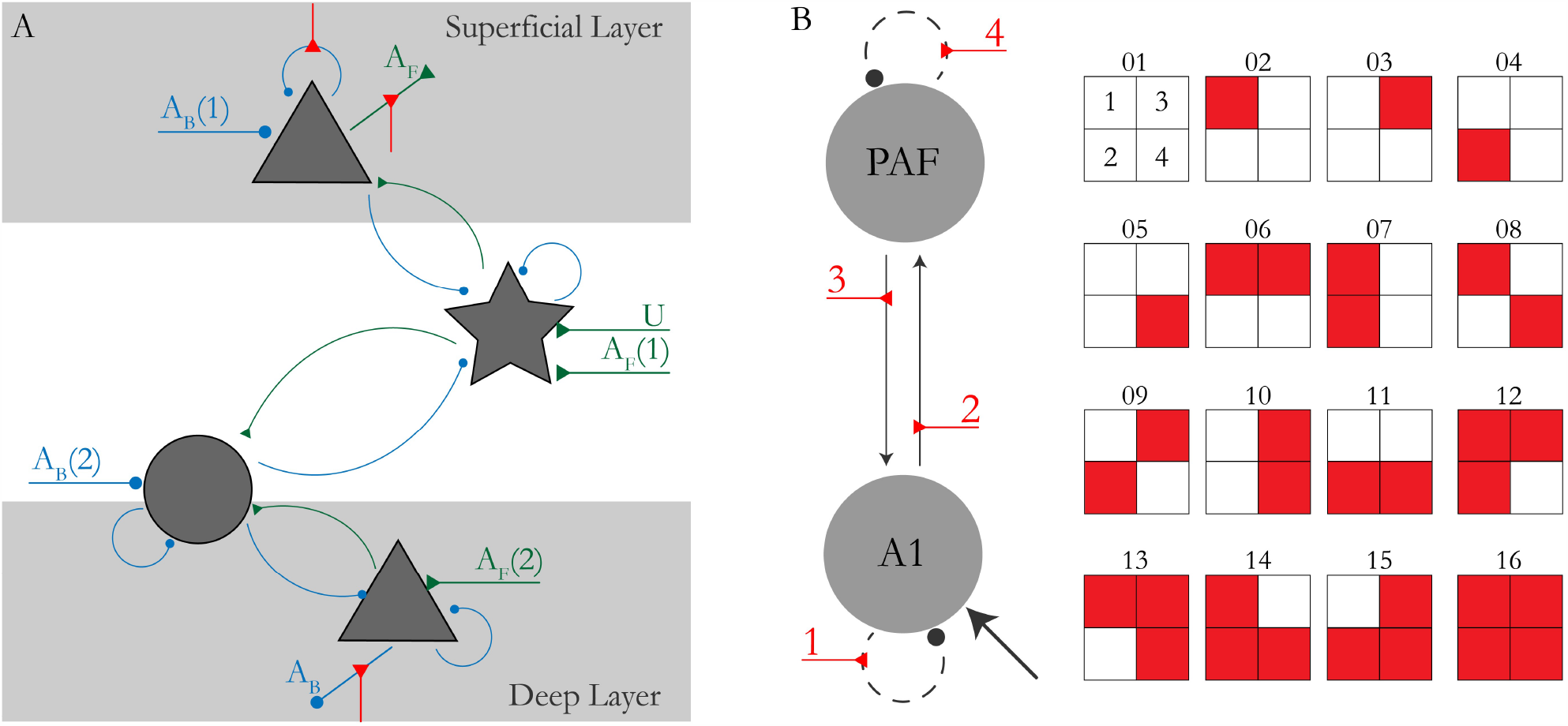
A) Connectivity pattern of the canonical microcircuit. Curved arrows indicate intrinsic (within region), straight arrows extrinsic (between region) or driving connections. Green colour indicates excitatory (triangular arrowheads), blue colour inhibitory connections (round arrowheads). Labelling indicates forward (A_F_), backward (A_B_) connections, or driving input (U) which entered A1. The index in the bracket refers to the corresponding parameter in the A-Matrix. Arrowheads show the direction of the connection. Red connections depict putative modulation by TONE (shown on the outgoing connections). Pyramidal cell populations are depicted by triangles, stellate cells by a star and inhibitory neurons by circles. B) Definition of model space. We consider 16 models, where connection strength (or excitability) can change by TONE. Red boxes indicate modulation by TONE, boxes 1 - 4 correspond to the connections on the left.

In our setting, the DCMs comprised two reciprocally connected sources, A1 and PAF. Driving input encoding auditory stimulation (by any tone) targeted region A1. Based on this basic structure, we explored a full factorial model space comprising all possible combinations of modulation by TONE (deviant vs. standard) on the forward connection, the backward connection and the intrinsic connection in both regions. This resulted in 2 = 16 models (Figure 1B), with 26 to 30 parameters (depending on the modulation structure).

We made a number of changes to the default implementation of the CMC in SPM12, motivated by prior testing of the framework on the non-pharmacological dataset. These changes included the use of a custom-written integration scheme for delay differential equations based on a continuous extension of Euler’s method for ordinary differential equations (Feldstein and Goodman, 1973; Schöbi, 2020). This was in response to questions about whether the default integration methods implemented in SPM12 are ideally suited for accurate parameter estimation (Lemarechal et al., 2018). Furthermore, default priors for the main analysis of the drug data were replaced by empirical priors from the non-pharmacological data (see paragraph on non-pharmacological dataset). Finally, in order to avoid local extrema, we ran the Variational Bayes inversion routine (i.e., Variational Laplace) under a multi-start approach by sampling 100 starting values from the prior.

### Model Selection and Averaging

Model goodness was assessed in terms of the negative free energy, which provides a lower-bound approximation to the log model evidence (Friston et al., 2007). We used random effects Bayesian Model Selection (BMS) (Stephan et al., 2009b) to compute the posterior probability that a specific model generated the data of a randomly selected subject from the group. Specifically, we computed protected exceedance probabilities (Rigoux et al., 2014) to compare models. This metric rests on the posterior probability *φ* that model *k* ∈ {1, …, *K*} is more probable than any other model *φ*_k_ = *p*(*r*_k_ > *r*_\k_|*y*) given the the group data *y*, while taking into account that differences may have arisen simply by chance.

Our primary interest, however, concerned the potential representation of drug effects in the estimated model parameters. Bayesian Model Averages (BMA) were calculated on the individual animal level to marginalize out model uncertainty (Penny et al., 2010). In other words, for a given model parameter, BMA computes its average posterior distribution over all models considered, where this average is weighted by the posterior model probabilities. We used the BMA estimates in all subsequent statistical tests.

### Statistics and Classification

Statistical analyses of the drug effects focused on estimates of DCM parameters that have a biological interpretation in terms of synaptic efficacy and plasticity. These include the connectivity (4), the kernel gain (6) and decay (4), and the modulatory parameters encoding a difference in response to deviant tones (4) (18 parameter estimates in total). For these parameter estimates, three different approaches were considered to test for pharmacological effects.

First, we computed a generic 1×5 ANOVA with a fixed factor DRUG and a random effect ANIMAL. We performed this test separately for each BMA parameter estimate as a dependent variable and used Bonferroni correction to correct for multiple comparisons.

Second, we tested for a parametric drug-effect relationship, where we use the notion of a ‘drug-effect’ as the change in parameter estimates, as we move from the drug with the most antagonistic effect (2 mg/kg scopolamine) to the drug with the most agonistic effect (6 mg/kg pilocarpine). This can be regarded as a more refined version of the ANOVA above, that leverages knowledge about generic dose-effect relationships. For this analysis, we specified a mixed effects model for the same estimates used in the generic ANOVA, assuming a linear fixed effect of DRUG, *X*_*drug*_ = […, −2, −1, 0 1, 2, …]^*T*^ corresponding to 1 and 2 mg/kg of scopolamine, vehicle, and 3 and 6 mg/kg of pilocarpine respectively and a random effect of ANIMAL (hemisphere specific). The dots indicate different rats/hemispheres. The corresponding general linear model is:

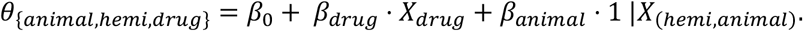

For each parameter vector *θ* (e.g. modulation of the forward connection), the values are ordered according to the subscript, i.e. animal, hemisphere and drug. Thus, *X*_*drug*_ codes for a linear effect over pharmacological interventions.

Finally, we tested for a mapping between the drug treatments and DCM parameter estimates using a linear support vector machine (SVM) with leave-one-out cross-validation (LOOCV) (Allwein et al., 2000). This procedure asks whether one could predict the drug label from the model parameter estimates. Specifically, this was based on the same 18 BMA estimates used in the classical tests above. Hyperparameters of the SVM were optimized within each cross-validation set (using nested cross-validation). In order to quantify the information that could be gained from the DCM parameters, we tested five different classifications: In four binary classifications, we compared the two extreme drug conditions against each other and individually against the vehicle condition, and the two antagonists vs. the two agonists. Finally, we applied a multiclass classification for all levels of the pharmacological factor. In order to be able to perform LOOCV in a balanced way, rats with data from only one hemisphere were omitted during classification. Please note that the left-out animal of the outer loop was not used in the inner loop, so no leaking of information was possible between test data and any parameter optimized within the inner loop. Importantly, in the LOOCV approach we adopted, it is not possible for the classifier to learn animal specific parameters based on the other hemisphere (which could be interpreted as a leak of information from the parameters of the other hemisphere of the same animal, or any other pharmacological condition).

To estimate the requisite confidence intervals, we used 1000 random permutations of the drug labels (within-animal and hemisphere), and re-estimated the full classification (including the optimization of the hyperparameters). The p-values then correspond to the proportion of permuted drug-label configurations that would have afforded a better classification than the actual one.

## Results

### Classical Analysis

From the 20 recorded hemispheres (10 animals), the data from three animals were excluded because of poor recording quality in either left and/or right hemisphere, based on visual inspection. For the remaining 14 hemispheres, all 5 pharmacological conditions were included in the analysis, resulting in 70 data points for statistical analysis.

First, we ran a mixed effects ANOVA with fixed effects TONE and PHARMA, their interaction and a random effect of ANIMAL. We found prolonged effects of TONE, PHARMA, and their interaction (see Figure 2) on evoked responses. These effects are consistent over the time window of interest, electrodes and deviant probability. The effect of TONE, evinced by the difference wave in Figure 2, exhibits two main peaks: an early negative peak around 25-50 ms and a “late” positive peak around 75-125 ms. It is also these two peaks that showed consistent interactions between TONE and PHARMA in all regions. Interestingly, the earlier peak is very dominant in the raw ERPs of both standard and deviant tones, most notably visible in the right hemisphere electrodes, with the two pilocarpine conditions exhibiting an additional dip right after 50 ms (see the arrow in Figure 2).

**Figure 2:**
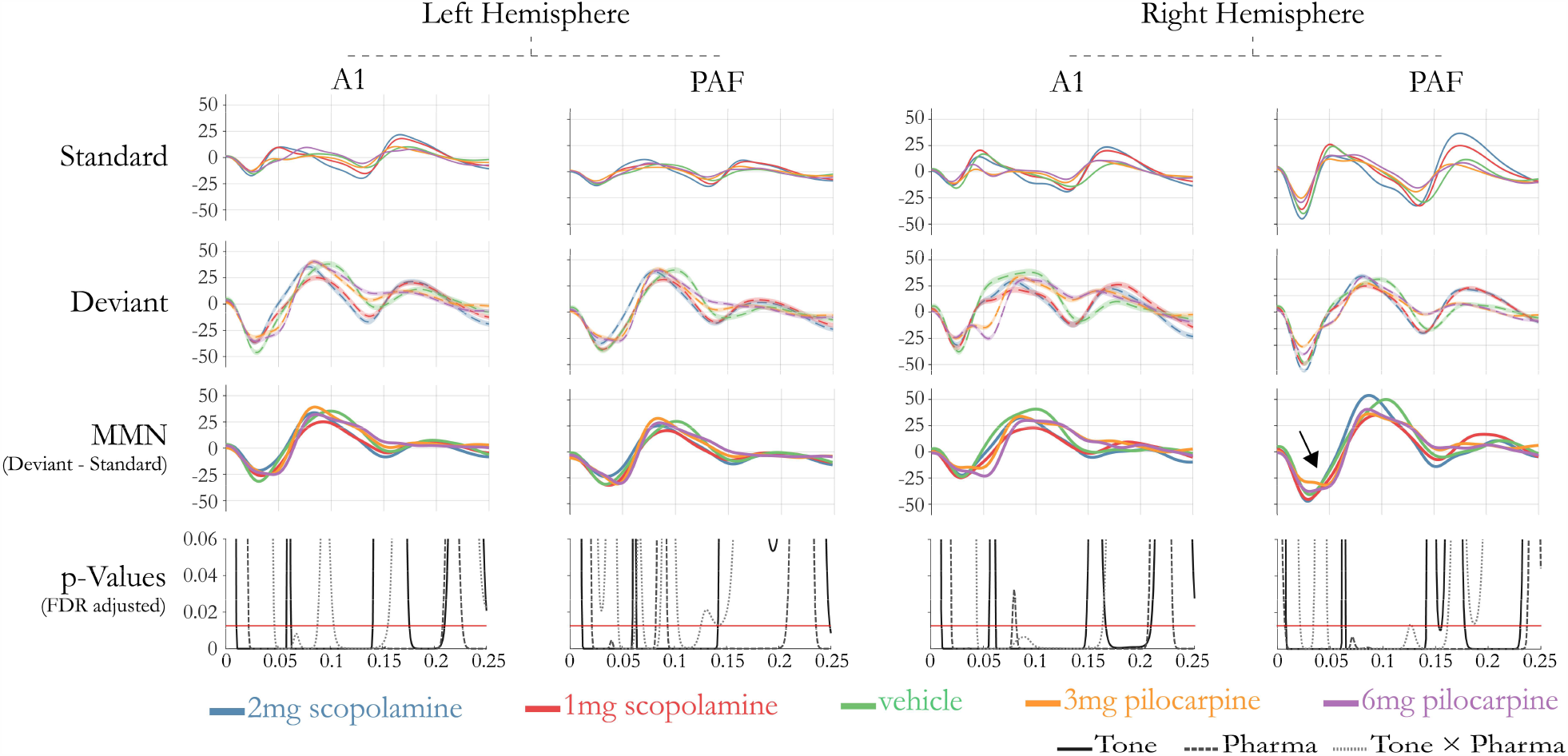
Grand Average Evoked Responses and results from the mixed effects ANOVA. Average (over animals and trials) Standard and Deviant tones are shown together with average difference waves for all drugs and both hemispheres (A and B). Statistical results show p-values (FDR corrected) of the main and interaction effects (without correction for electrodes). The red line indicates a threshold of p = 0.0125 (Bonferroni correction over four electrodes). The arrow indicates the additional dip for the two agonistic interventions mentioned in the paragraph.

### Dynamic Causal Modelling

The animal-specific ERPs were modelled using DCM. Notably, for consistency with the classification results described below, we use only those rats where both hemispheres were included in the data analysis (N=7).

Using multi-start Variational Laplace, we inverted each of the 16 models shown in Figure 1 initializing the gradient ascent with 100 different starting values (sampled from the prior over parameters), for each rat, pharmacological condition, and hemisphere.

In terms of the primary measure of model goodness – the (negative) free energy – the multi-start approach was clearly beneficial. The model inversion with the highest free energy estimate was always from a starting point that was not the prior mean of the parameters. Starting from the prior mean is the default often used. This finding illustrates the multi-modal nature of the objective function and the utility of multi-start procedures.

Random effects model selection between the 16 competing DCMs did not yield a conclusive result (Figure 3A), although there was a tendency for more complex models to perform better, especially for the agonist conditions where the (protected) exceedance probability was approaching 0.95 (Rigoux et al., 2014). The most complex model (model 16) performed consistently well across all pharmacological conditions. Runner ups included models of greater complexity, such as models 7, 9, 11, 12 and 15. Common to all these models is the presence of a modulation of the forward connection.

**Figure 3:**
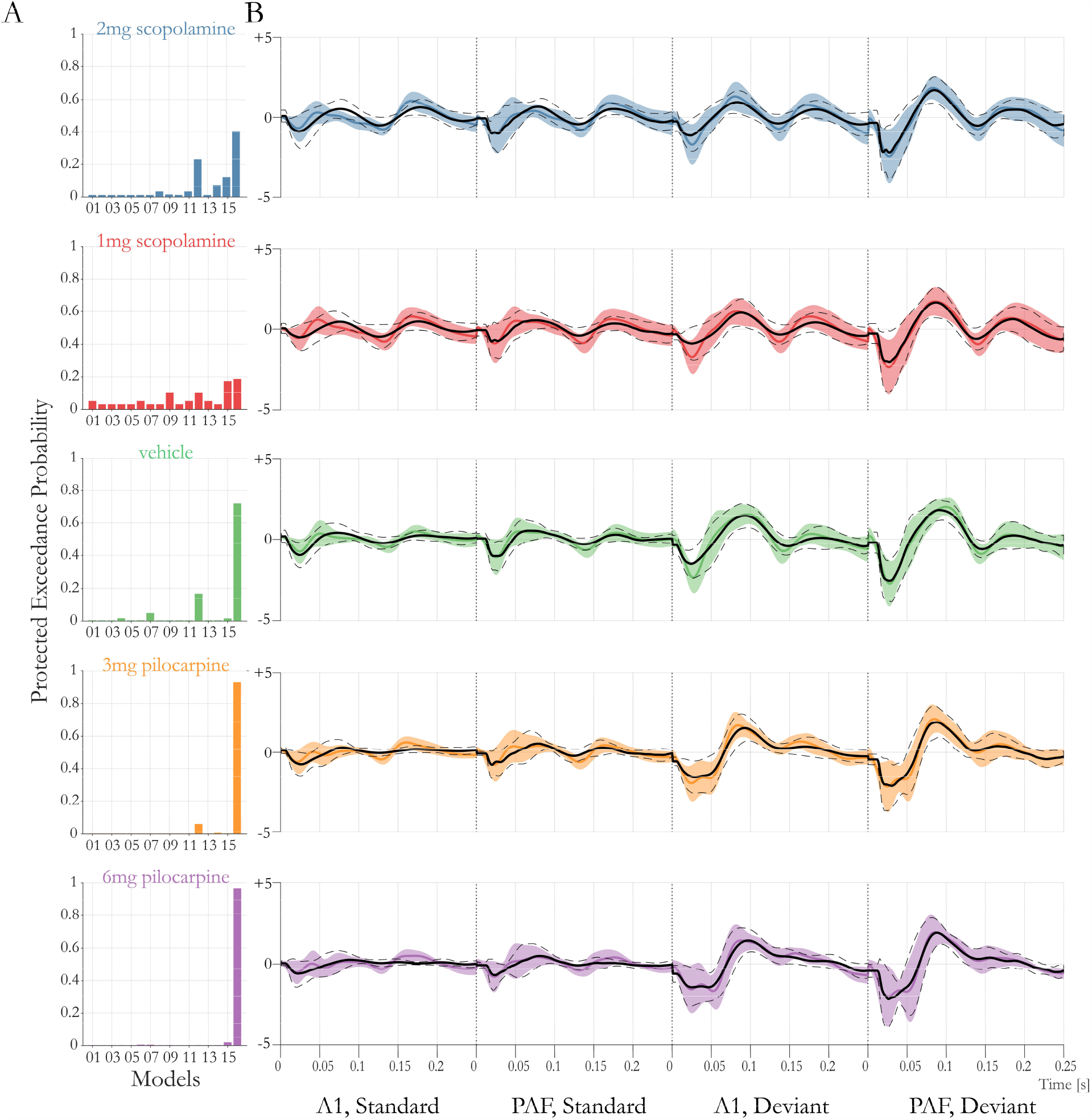
A) Bayesian Model Selection (BMS). Protected Exceedance probabilities reported for all sixteen models and drugs. B) Average (over animals) data (coloured line) and prediction (black solid line) for model 16. Shaded area depicts standard deviation of the data (over animals and hemispheres), dotted lines depict standard deviation of prediction (over animal and hemispheres).

The overall fit for the winning (most complex) model is illustrated in Figure 3B by comparing the average prediction of the model (averaged over both hemispheres and rats) and empirical data. For reference, the average prediction would explain 88 % (2mg scopolamine), 88 % (1mg scopolamine), 93 % (vehicle), 93 % (3mg pilocarpine), 93 % (6mg pilocarpine) of the average signal variance.

### Parameter Estimation and Statistics

Since there was no unambiguous winning model in all pharmacological conditions, we computed BMAs on the individual animal level, effectively marginalizing out the model from the posterior distributions. Our primary interest were parameters with a biological interpretation in terms of synaptic processes, i.e., extrinsic connection strengths, modulatory influences, kernel gains and decays (in total 18). We used these BMA estimates in two separate ANOVAs. First, we tested for any effect of DRUG, while correcting for the random effect of ANIMAL. Second, we tested for a linear effect of drug (i.e. across the different levels of muscarinic effects, from the highest antagonistic via vehicle to the highest agonistic dose). ANOVAs were computed for each parameter of interest and Bonferroni corrected for the ensuing 18 tests. The results are summarized in Table 1 and Figure 4. For the one-way ANOVA with random effect ANIMAL, there was a significant effect on the kernel gain of self-inhibition of the superficial pyramidal cell in PAF, and on the kernel decay of the inhibitory cell, *p* < 0.05 (corrected). The latter parameter is set to be the same for both regions. When testing for a linear effect of drug, we observed a significant linear relationship in five parameters: The forward connection to the deep pyramidal layer (c.f. A_F_(2) in Figure 1), the modulation of the forward connections (A1→PAF), the modulation of the backward connections (PAF→A1), and the same two kernel parameters found in the previous ANOVA.

**Table 1:** ANOVA statistics on the BMA estimates for the classical ANOVA, and the ANOVA where DRUG was treated as a factor with a linear effect from the most antagonistic to the most agonistic drug condition.

|  |  |  | classical |  | linear |  |
| --- | --- | --- | --- | --- | --- | --- |
| Class | Connection | Parameter | F-Values | p-Values<br>(uncorrected) | F-Values | p-Values<br>(uncorrected) |
| A Matrix<br>(A) | SPC → SC | $A_F(1)$ | 1.3705 | 0.2551 | 5.3372 | 0.0242 |
| | SPC → DPC | $A_F(2)$ | 2.5341 | 0.0496 | **10.1793** | **0.0022 (-)** |
| | DPC → SPC | $A_B(1)$ | 1.0172 | 0.406 | 0.4711 | 0.4950 |
| | DPC → IC | $A_B(2)$ | 0.6058 | 0.6598 | 0.9121 | 0.3429 |
| Modulation<br>(B) | $A1 \rightarrow A1$ | B1 | 2.8086 | 0.0334 | 5.8079 | 0.0189 |
| | $A1 \rightarrow PAF$ | B2 | 3.8664 | 0.0074 | **12.455** | **0.0008 (+)** |
| | $PAF \rightarrow A1$ | B3 | 4.2405 | 0.0044 | **16.8168** | **0.0001 (+)** |
| | $PAF \rightarrow PAF$ | B4 | 0.5758 | 0.6813 | 0.0043 | 0.9479 |
| Kernel<br>Gain<br>(G) | SPC → SPC (A1) | G1 | 1.1165 | 0.3564 | 1.9527 | 0.1668 |
| | SPC → SPC (PAF) | G2 | **4.6626** | **0.0023** | **17.9828** | **0.0001 (+)** |
| | SPC → SC (A1) | G3 | 0.8903 | 0.4755 | 1.5026 | 0.2249 |
| | SPC → SC (PAF) | G4 | 1.3646 | 0.2572 | 0.6301 | 0.4303 |
| | IC → SC (A1) | G5 | 1.5008 | 0.2123 | 4.8325 | 0.0313 |
| | IC → SC (PAF) | G6 | 2.9784 | 0.0262 | 8.6147 | 0.0047 |
| Kernel<br>Decay<br>(T) | SC | T1 | 0.7326 | 0.5734 | 0.793 | 0.3766 |
|  | SPC | T2 | 0.7238 | 0.5792 | 1.3634 | 0.2474 |
|  | IC | T3 | **4.62** | **0.0026** | **16.2289** | **0.0002 (-)** |
|  | DPC | T4 | 1.6404 | 0.1761 | 4.0477 | 0.0486 |

**Figure 4:**
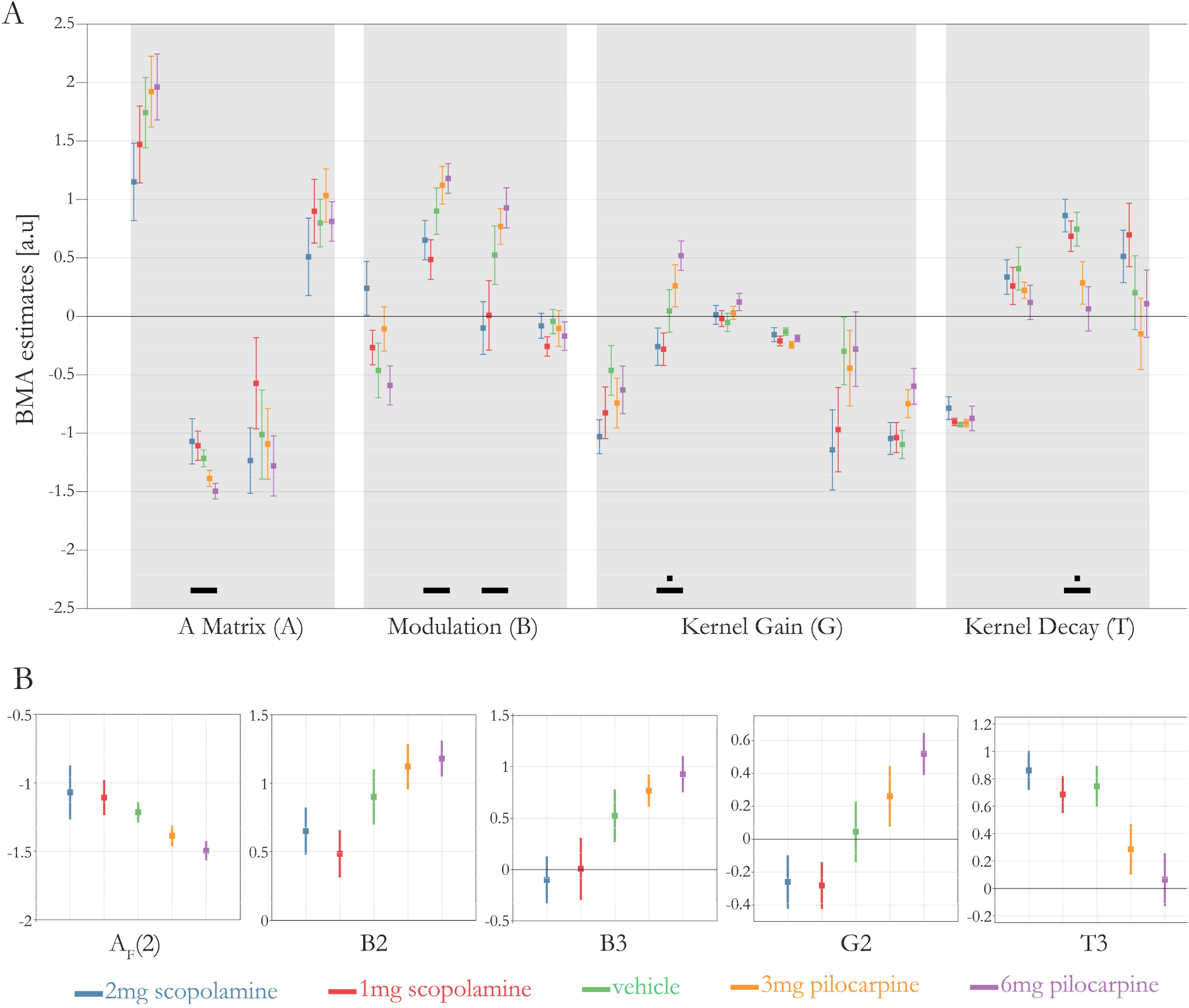
A) BMA estimates for all animals (n=7). BMAs are computed on the first level and pooled over both hemispheres. Error bars depict SEM. Mixed effects (MFX) ANOVA on the BMA parameters are displayed. We considered two MFX models. First, a model with fixed factor DRUG (5 levels) and random effect ANIMAL. Black squares indicate significant results at p < 0.05 (Bonferroni corrected). Second, a MFX ANOVA with a linear, fixed effect of DRUG and random effect of ANIMAL. Black horizontal bars indicate significant results at p < 0.05 (Bonferroni corrected). B) Zoomed in display of parameters showing a significant linear effect. Labelling according to Table 1.

### Classification

Finally, we asked whether it was possible to predict the drug label (or even level) from the model parameter estimates. We used the BMA estimates as features for a linear SVM with LOOCV. Here, in each fold, the classifier was trained on the drug labels of all but one rat and then the drug labels of the left-out rat was predicted. We computed the balanced accuracy (BA) as performance score of classification and considered the five classifications described in the Methods. We were able to predict the individual drug levels in a multiclass classification with 31.4% BA (*p* = 0.024, chance level: 20%). Also, we could distinguish between the most extreme antagonistic and agonistic effects with 92.9% BA (*p* < 0.001, chance: 50%), between the highest dosage of pilocarpine and vehicle with 71.4% BA (*p* = 0.032, chance: 50%), and between both drugs with antagonistic and agonistic effects with 73.2% BA (*p* = 0.001, chance: 50%). Classification between the highest dosage of scopolamine and vehicle was not significantly different from chance, with 39.29% BA (*p* > 0.10. Classification results are summarized in Figure 5. All *p*-values reported here are based on permutation test on the drug labels and were not corrected for multiple comparisons. However, all classifications with *p* < 0.01 are significant when Bonferroni corrected for the five tests under the chosen alpha level (indicated by two stars in Figure 5F). Note that the permutation based statistical testing of classification accuracies works robustly even in the context of relatively few data points (Varoquaux et al., 2017), as is the case here.

**Figure 5:**
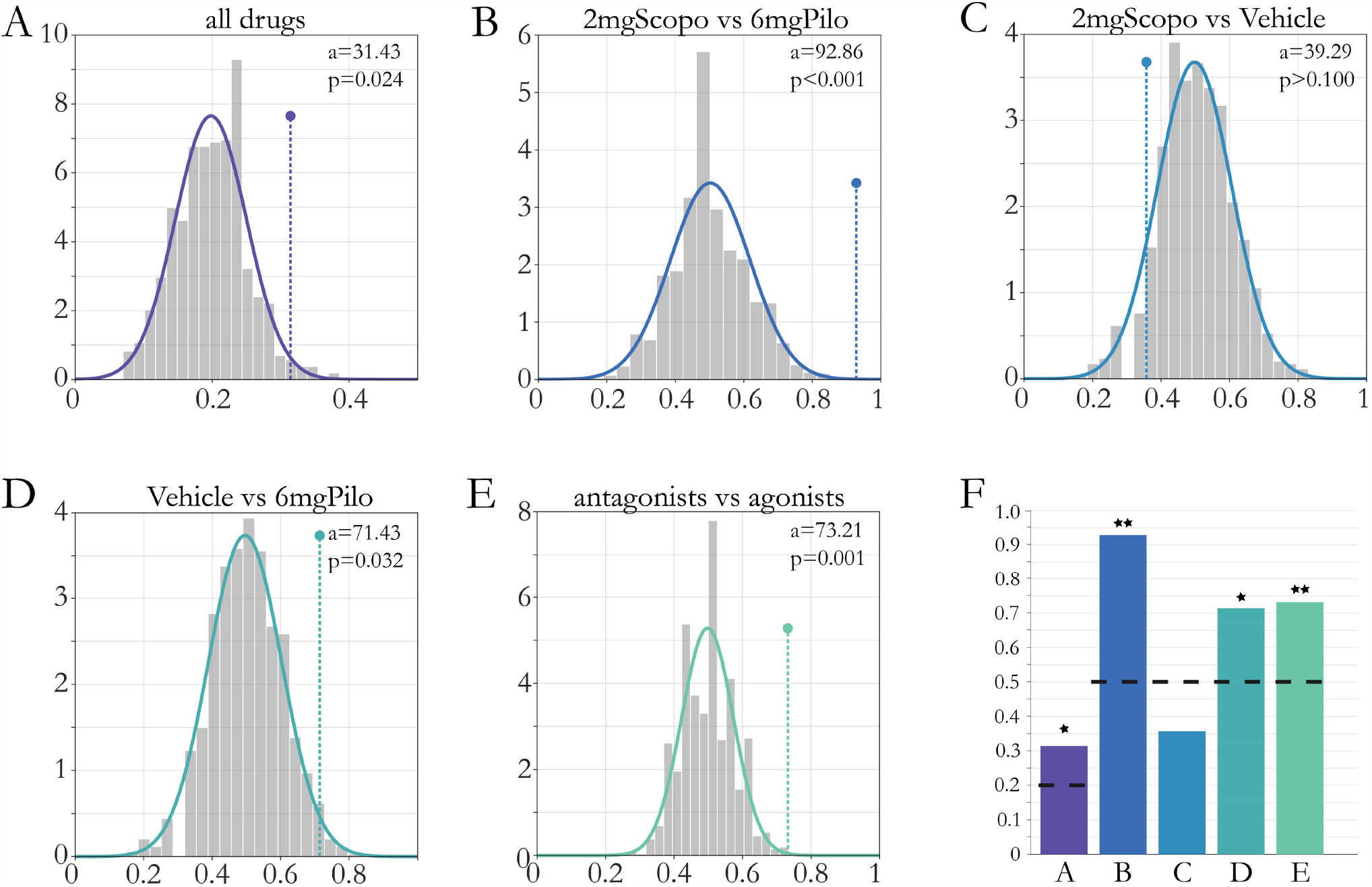
Permutation statistics for multiclass (A) and binary (B - E) classifications on the BMA results. Grey bars depict cross validation (CV) accuracies of permuted labels, solid line a gaussian fit on the histogram. Dotted line depicts CV accuracy for the true labels. Numbers refer to the CV accuracy of the true labels (a) and the percentage of a permutations leading to a higher accuracy (p). F) Balanced Accuracies for all classifications in (A-E). Stars indicate significance at p<0.05 (1 Star) and p < 0.01 (2 Stars) based on permutation statistics. The black dotted line indicates chance level for the specific classifications.

## Discussion

In this study, we investigated changes in epidural EEG recordings induced by graded pharmacological manipulations of muscarinic receptors during the auditory MMN. Using physiologically interpretable DCMs of auditory circuits, we were able to explain ERP changes across different levels of muscarinic receptor function in terms of synaptic connections likely affected by the drugs. We then identified several model parameter estimates that exhibited significant linear drug-effect relationships. Finally, we demonstrated that the estimated synaptic parameters allowed us to predict drug type (antagonist versus agonist) with nearly 93% accuracy and, less precisely, the dose under which a given dataset had been recorded. To our knowledge, this represents the first study using a graded manipulation of a neuromodulatory transmitter during the auditory MMN, from strong/weak inhibition via placebo to weak/strong enhancement, using highly selective drugs and multiple recordings from both hemispheres in awake rodents.

### Generative embedding – using DCM for model-based feature reduction

When trying to predict the pharmacological state from the recordings, one faces the problem that the dimensionality of the feature space (the recordings over time) is far greater than the number of labels (drug status) to be predicted. A useful approach to address this problem is generative embedding (Brodersen et al. 2011). Simply speaking, generative embedding uses a generative model to partition the data into unexplained signal (noise) and explained signal and uses the parameter estimates as a compact, low-dimensional representation of the latter. That is, instead of using a noisy, high-dimensional set of features (the original recordings), generative embedding uses a low-dimensional and de-noised feature set for subsequent (un)supervised learning. While the choice of a specific generative model depends on the data type and the modeler’s theory about the processes that generated the data, DCM has been a frequent and successful choice in previous generative embedding studies of brain activity measurements (Brodersen et al., 2011; Brodersen et al., 2014; Frässle et al., 2020).

In the present study, Dynamic Causal Modelling allowed us to reduce the dimensionality of the problem from 1000 features (datapoints per recording) to 30 (or 18) features (parameter estimates), while maintaining the overall information contained in the complete time series (as demonstrated by the good model fits). Notably, when performing generative embedding, it is not necessarily the goal to fit each specific (traditional) feature of classical ERP analyses, such as amplitudes or latencies of MMN components, but to reproduce the overall data as the output of a physiological system, reducing the dimensionality to some key parameters. As a concrete example, let us consider the first peaks (around 25 ms and 50 ms) of the primary auditory region for the 1mg scopolamine conditions (see Figure 3B): The timing of the second peak is not perfectly captured by the model. In order to fit this specific feature, one would have to adapt the model, e.g. by reducing constraints, adding degrees of freedom, or using tailored noise functions that force the algorithm to fit well in certain temporal windows. However, such changes that are tailored to explain specific features of the data would increase the risk of overfitting in general (with detrimental effects for subsequent out-of-sample classification based on the parameter estimates) and are not in line with the goal of the present study. Here, the emphasis was on obtaining a plausible but compact representation of the overall signal and how it is affected by drugs. The fact that our model fits result in high proportions of variance explained across all pharmacological conditions indicate that information of the global signal was adequately captured by the model.

### Effects of auditory deviance

Model comparison suggested that both forward and backward connections within a small auditory circuit comprising primary (A1) and secondary (PAF) areas were modulated by the occurrence of surprising tones (deviants). This fits well with predictive coding accounts of the MMN, where surprising events lead to (precision-weighted) prediction error updates of an internal model, in order to minimize surprise (Baldeweg, 2007; Garrido et al., 2008; Garrido et al., 2009). More specifically, previous modelling studies of the MMN suggested that the occurrence of deviants modulate long-range glutamatergic connections as well as local gain adaption (Garrido et al. 2008; Moran et al. 2013). Our results are consistent with these findings, with slightly reduced emphasis on local gain modulation.

### Drug-effect relationships

In addition to the classical ANOVA, the modulation of the A1→PAF forward connection exhibited a linear drug-effect relationship. The backward modulation also showed a significant positive linear relation to drug level. In other words, the more strongly muscarinic receptors were activated, the stronger the increase in forward and backward connection for deviant tones. This finding is in contrast with a previous study in humans which investigated the effect of galantamine on the auditory MMN and reported mainly local gain increases in A1 (Moran et al., 2013). It is possible that this difference is due to the different action of galantamine which increases the level of ACh in general and may also allosterically potentiate nicotinic receptors ((Samochocki et al., 2003); but see (Kowal et al., 2018)). Physiologically, the drug-induced changes in glutamatergic long-range connections between auditory areas (which presumably draw on both AMPA and NMDA receptors; see discussion in (Schmidt et al., 2012)) could be mediated by short-term changes in synaptic transmission. Specifically, muscarinic agents are known to change AMPA and NMDA receptor function by various mechanisms, including phosphorylation or changes in subunit composition (Marino et al., 1998; Grishin et al., 2005; Shinoe et al., 2005; Di Maio et al., 2011; Lopes et al., 2013; Zhao et al., 2018; Zhao et al., 2019), for review, see (Butcher et al., 2009).

Linear but not deviant-specific pharmacological effects were found in two of the DCM parameters. To discuss these results in more detail, we consider their effects on the two pyramidal cell (PC) populations (see Figure 1), since those directly contribute to the measured EEG signal. We observed an increase in the gain of inhibitory self-connections of superficial PCs in the PAF, resulting in a smaller (in absolute values) initial peak of the ERP. Furthermore, there was a decrease in the time constant of the inhibitory cell, i.e. faster decay. The inhibitory cell directly drives deep PCs but has no direct influence on superficial PCs. Since deep PCs are (intrinsically) driven only by ICs in the model, a faster decay of inhibition will result in less deactivation of deep PCs. This, in turn causes the overall signal to decay more slowly after the first peak. Both results – a graded expression of (absolute) amplitude and its decay back to zero – can be observed in the ERPs in Figure 2, around 25-50 ms. A similar dichotomy of muscarinic action into a fast (net inhibitory) and a slower depolarizing effect was observed in vitro (McCormick and Prince, 1985).

A central aim of the present study was to test the feasibility of predicting muscarinic receptor status, out of sample and from parameter estimates of a physiologically interpretable circuit model. The strategy of using parameter estimates from a generative model for subsequent (un-) supervised learning is known as ‘generative embedding’ (Brodersen et al., 2011) and plays a central role in attempts to establish computational assays for psychiatry (Stephan et al., 2017). For example, recent work suggested that dopaminergic and cholinergic alterations can be predicted out-of-sample from eye movements (Aponte et al., 2020). The generative embedding approach has two main advantages: it offers a theory-led dimensionality reduction (from high-dimensional noisy data to a small set of model parameter estimates), and it enables the interpretation of machine learning results in terms of biological mechanisms represented by a model.

In this study, generative embedding suggested that a relatively simple model of a small cortical circuit can be used to predict muscarinic receptor status from EEG data. When considering the different pharmacological conditions separately, the most robust discrimination was obtained under the muscarinic agonist pilocarpine. That is, all classifications involving pilocarpine (Figure 5A, B, D and E) resulted in balanced accuracies significantly above chance, and the higher the difference in dosage, the better the classification. By contrast, distinguishing the muscarinic antagonist scopolamine from placebo proved more challenging. There could be several reasons for this, including drug differences of neuronally effective dosage regimes or strong non-linearities in drug-effect relationships.

While the classification accuracies for different dose levels are not yet close to clinically required levels of precision, the more general question of whether muscarinic receptor function had been diminished or enhanced (antagonist vs. agonist) could be answered decisively with a balanced accuracy above 90%. If this result could be replicated in a human EEG study – with a sufficiently large sample size – a computational assay for distinguishing hyper-vs. hypo-activity of muscarinic receptors might be feasible. As described in the Introduction: given the importance of individual neuromodulatory differences in schizophrenia (Stephan et al 2009), the likely existence of schizophrenia subgroups with differences in muscarinic receptors (Scarr et al., 2009; Scarr et al., 2018), and the distinctive anti-muscarinic properties of clozapine and olanzapine as two of the most potent antipsychotics (for review, see (Kapur and Remington, 2001), such an assay could find important clinical applications for differential diagnosis and treatment selection in schizophrenia.

### Limitations and outlook

While relatively large for rodent studies with in vivo recordings, the sample size of our study is not sufficient to quantify out-of-sample prediction accuracy with high precision (Varoquaux, 2018). However, even in small sample scenarios, permutation tests (as used in our study) yield a robust and valid measure of whether the estimated prediction accuracy is significantly different from chance (Varoquaux et al., 2017). Still, the results should be interpreted with caution and need to be replicated in (human) studies of larger size.

Another limitation is that the generative model used in this study does not allow one to directly map synaptic parameters onto a particular neurotransmitter system. In other words, there is no single parameter in our model that explicitly represents muscarinic function. Instead, it is likely that we are observing a net effect of pharmacologically altered muscarinic receptor function on several mechanisms represented in the model, like synaptic connectivity strength and neuronal gain. For example, it is known that muscarinic receptors change glutamatergic synaptic transmission through influencing both NMDA and AMPA receptors (Marino et al., 1998; Grishin et al., 2005; Shinoe et al., 2005; Di Maio et al., 2011; Lopes et al., 2013; Zhao et al., 2018; Zhao et al., 2019); an effect that can be (and was) observed in the estimates of model parameters encoding glutamatergic long-range connections. Similarly, muscarinic receptor activation strongly affects neuronal excitability and gain (McCormick et al., 1993; Shimegi et al., 2016); this effect is captured by estimates of parameters representing the gain of postsynaptic kernels. Here, however, our model suggests an increase in self-inhibition of the superficial PC population, which differs from reports of muscarinic agents reducing GABA release (Salgado et al., 2007). Interestingly, the majority of DCM studies on cholinergic modulation or non-pharmacological interventions thought to affect neuromodulation (e.g. attention and gain control) identify inhibitory connections involving superficial PCs; especially those trying to explain high-frequency (gamma) induced responses (Fogelson et al., 2014; Pinotsis et al., 2014; Auksztulewicz and Friston, 2015; Bastos et al., 2015; Shaw et al., 2017).

Our findings suggest that using DCM for model-based feature extraction is a promising way forward in order to classify pharmacological status based on ERP data. We emphasise that generative embedding is not the only option for this classification attempt. One could in principle use any other feature derived from the EEG time series (e.g. specific ERP features) or even the raw time series as input to the classifier. Generally, there is a huge number of options how features for classification could be extracted from the rich dataset at our disposal. However, a comparison of different feature sets was deliberately not attempted as it was outside the scope of our study (see the a priori analysis plan at https://gitlab.ethz.ch/tnu/analysis-plans/schoebietal_auditory_mmn_dcm_2020). In this study, the goal was not to claim any superiority of generative embedding compared to other possible approaches. Instead, we set out to test whether parameter estimates of a specific model (DCM) are plausibly linked to pharmacological manipulations and can afford predictions of drug status.

To date, no generative models exist that represent neuromodulatory transmitter action directly, through distinct biophysical parameters (although see (Fogelson et al., 2014) and (Auksztulewicz and Friston, 2015) for an application of DCM equipped with top-down neuromodulatory connections). This represents an area of active ongoing research. Our current findings – that changes in muscarinic receptor function can be inferred from auditory mismatch signals using a generative model (DCM) of a cortical circuit – illustrate the potential of generative embedding for developing computational assays for psychiatry. Pending further improvements and validation in human populations, such assays might eventually play a useful role for differential diagnosis, stratification, and treatment predictions in heterogeneous psychiatric disorders.

## Acknowledgements

This work was supported by the René and Susanne Braginsky Foundation (KES), the University of Zurich (KES) and the Max Planck Society (MT). KJF was funded by a Wellcome Trust Principal Research Fellowship (Ref: 088130/Z/09/Z)

## Footnotes

1 The all-caps notation is used to explicitly refer to the factor in the statistical model.

